# Development of an Ultra Low-Cost SSVEP-based BCI Device for Real-Time On-Device Decoding

**DOI:** 10.1101/2022.01.29.478203

**Authors:** James Teversham, Steven S. Wong, Bryan Hsieh, Adrien Rapeaux, Francesca Troiani, Oscar Savolainen, Zheng Zhang, Michal Maslik, Timothy G. Constandinou

**Affiliations:** Department of Electrical and Electronic Engineering and Centre for Bio-Inspired Technology, Imperial College London, South Kensington Campus, London SW7 2AZ. UK; Department of Life Sciences, Imperial College London, South Kensington Campus, London SW7 2AZ. UK; UK Dementia Research Institute (UKDRI), Care Research and Technology (CR&T) Centre at Imperial College London and the University of Surrey

## Abstract

This study details the development of a novel, approx. £20 electroencephalogram (EEG)-based brain-computer interface (BCI) intended to offer a financially and operationally accessible device that can be deployed on a mass scale to facilitate education and public engagement in the domain of EEG sensing and neurotechnologies. Real-time decoding of steady-state visual evoked potentials (SSVEPs) is achieved using variations of the widely-used canonical correlation analysis (CCA) algorithm: multi-set CCA and generalised CCA. All BCI functionality is executed on board an inexpensive ESP32 microcontroller. SSVEP decoding accuracy of 95.56 ± 3.74% with an ITR of 102 bits/min was achieved with modest calibration.

## I. Introduction

Commercial EEG-based BCIs are typically priced in excess of several hundred GBP [1]–[3]. This makes them largely inaccessible for experimentation on a mass scale, especially outside well-funded academic settings. A low-cost, easy-to-use BCI device would likely prove useful for gathering EEG data in a more general population, as well as be an invaluable pedagogical tool for engaging and educating young learners about basic neurophysiology and neurotechnology; fields that would otherwise be difficult to access below a tertiary education level. The core objective of this study was to develop a mobile EEG-based BCI that is widely accessible both in terms of affordability and ease of use. With this objective in mind, the following constraints were devised:

1. A budget of ∼£20 per device.
2. Self-contained hardware with edge computing.
3. Minimal user calibration and no specialised equipment.
4. Real-time decoding and feedback of signals.
5. Device comfort: non-invasive and cannot use ‘wet’ electrodes that would hamper the user experience.

While several non-invasive EEG-based BCI paradigms exist, the steady-state visual evoked potential (SSVEP) has received much attention owing to the fact that it is easy to implement, requires no user training and can achieve relatively high information transfer rates (ITRs) [4]. Upon focusing on a flashing visual stimulus, SSVEPs are involuntary electrical potentials that are produced by the brain’s visual cortex at the same frequency of the stimulus and its harmonics [5]. By choosing which stimulus is observed, one produces SSVEPs that can be decoded. This form of BCI control was chosen for the project as it is highly mobile and does not require specialised lab equipment.

Many SSVEP decoding algorithms exist for this purpose and have been widely studied [6]–[9]. It is often reported that including historical ‘training’ or calibration data - whether subject specific or across multiple subjects - offers significantly superior decoding performance compared to inference on instantaneous test signals (as with standard CCA) [7]–[9]. However, the project constraints require that all decoding-related computation happen on device and that all the required data be stored in memory on hardware. As such, two extensions of the widely-used canonical correlation analysis (CCA) algorithm are explored in this study: multiset CCA (MsetCCA) [8] and generalised CCA (GCCA) [9]. Both algorithms were chosen for their computational tractability on the ESP32 microcontroller, superior efficacy and robustness over standard CCA due to the use of historical data, and their effectiveness even when using few calibration trials.

### A. Previous works

Several low-cost, lightweight BCI devices have been developed in the EEG research community. These consist of promising BCI devices that have demonstrated reliable EEG signal decoding in significantly resource-constrained settings [10]–[14]. In particular, these constraints are computing capacity, mobility or form factor, number of channels available and cost.

Uktveris et al. demonstrated a very promising device with wireless capabilities (communication with a host device using Bluetooth Low Energy (BLE) and/or Wi-Fi) and some on-device digital signal processing ability [11]. However, their device requires final signal decoding to be performed on a host device separate from the actual BCI itself. Furthermore, the cost of the raw materials in their project was around C114 [11], although their system can measure 16 to 64 channels depending on sampling rate. As such, it is a compact and low-power EEG recording device but did not seek to do the fully on-implant decoding that was the objective of this work.

The authors in [13] created an embedded BCI system that consists of a small, analogue sampling device that communicates using RF signals with an external controller. However, all processing and decoding occurs on the external decoder. Across three mental tasks, three SSVEP frequencies and eyes closed, they achieved an average classification accuracy of 74% and an average information transfer rate (ITR) of 27 bits/min.

Acampora et al. provide a single-channel EEG device in [15] and an expansion for SSVEP decoding in [14]. This device achieved a maximum SSVEP classification accuracy of 74.5% for a 2 s recording window, up to 92.7% over a 4 s window and 97.6% for a 6 s window for four SSVEP frequencies, which is state of the art. It used a support vector machine with Gaussian or RBF kernel. However, this device uses an off-the-shell Olimex EEG-SMT device for signal acquisition and a Raspberry Pi 3 for processing and decoding. While the device is effective and mobile, it significantly exceeds the £20 price budget in this project.

## II. Methods

### A. System architecture

Fig. 1 shows a block diagram of the system hardware that comprises a microcontroller (MCU) platform for data acquisition, processing & communication, a power management system, an analogue front-end for signal conditioning and a user interface with dry electrodes, two buttons and two LEDs. The electronics and PCB cost less than £10 each at 100 devices volume.

**Fig. 1:**
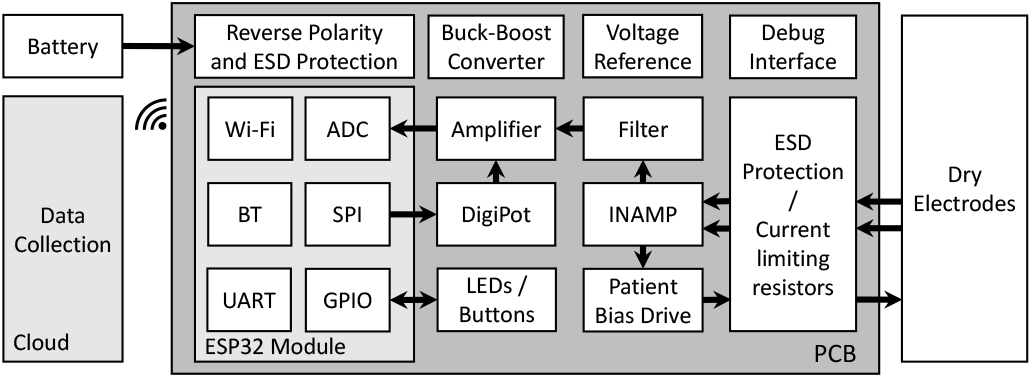
System block diagram

The system data flow is illustrated in Fig. 2. Acquired EEG signals are digitised, filtered, downsampled, and decoded on the device. The raw and decoded SSEVP signals are streamed wirelessly via MQTT to a cloud service for human-computer interaction.

**Fig. 2:**
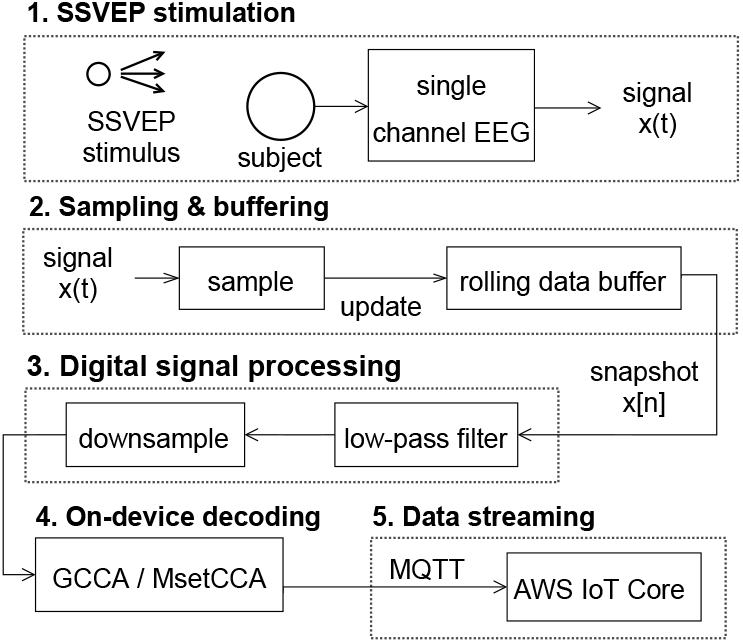
Flowchart showing system data flow

### B. Apparatus

A hardware break-down of the device shown in Fig. 3 is provided below.

**Fig. 3:**
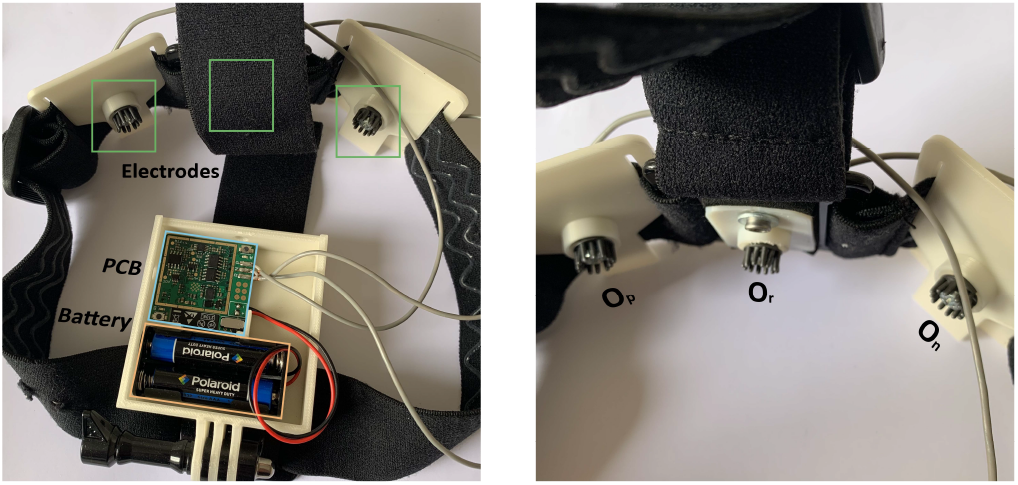
Images of the NGNI BCI device. Left: Adjustable elastic headband with fully-contained electronics. Right: Close up of the reference (centre) and two active electrodes.

#### 1) PCB

Central to PCB module is an Espressif ESP32 SoC based on the Tensilica Xtensa LX6 microcontroller featuring a dual-core 240MHz CPU. It supports Wi-Fi and Bluetooth connectivity, SPI, UART, and GPIOs. Following sampling at *f*_*s*_ = 256 Hz by the on-board 12-bit successive approximation (SAR) ADC, preliminary signal processing is performed on the analogue front-end module comprised of the following components: an instrumentation amplifier with common-mode feedback to the user as bias, a third-order Hourglass Sallen-Key filter with a 50 Hz notch and a 37.4 Hz corner frequency, and a digital potentiometer (DigiPot) based variable gain amplifier for fine tuning signal gain.

#### 2) Electrodes

The headset features a single reference electrode (O_r_) and two active electrodes (O_p_ and O_n_) from which a differential measurement is taken to yield a single channel EEG signal.

#### 3) Battery module

Multiple battery types are supported, including commercially available AAA/AA or single-cell lithium batteries. A buck-boost converter generates the digital supply voltage for the MCU module, and a voltage reference is used to supply the analogue front end circuitry and generate the half-rail voltage reference.

### C. Stimulus Design

The effect of SSVEP frequency and colour (among some other factors) on decoding accuracy and signal-to-noise ratio (SNR) was investigated in [16]. Of those tested, stimulus frequencies in the region of 12 Hz were found to maximise SNR [16]. Of colours tested, white was most effective around 12 Hz and green for lower frequencies towards 5 Hz. Red was found to increase risk of inducing epilepsy and so should be avoided [16]. Frequencies of around 12 Hz were also found to cause less visual fatigue than lower frequencies. Accordingly, stimulus frequencies of 7, 10 and 12 Hz were used in this study.

To allow for cross-platform, cross-device deployment, this interface was implemented as a lightweight HTML page with basic CSS styling and Javascript to handle animation (flickering of the squares).

### D. Digital Signal Processing

Due to embedded memory constraints and analogue filter dynamics, *N*_*h*_ = 1 harmonic was used for the harmonic reference signal where applicable. This implies a maximum signal frequency of 24 Hz (1^st^ harmonic of 12 Hz stimulus). Accordingly, a 10^th^ order elliptical low-pass filter with corner frequency *f*_*c*_ = 26 Hz was designed to meet a maximum pass-band ripple of *r*_*p*_ = 0.2 dB and a minimum stop-band attenuation of *r*_*s*_ = 80 dB. The filter was implemented in firmware as cascaded lower-order segments for improved numerical stability and less sensitivity to quantisation errors in filter coefficients. Following low-pass filtering for rejection of 50 Hz interference, signals are downsampled to 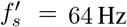 to reduce memory and computational demands while satisfying the Nyquist criterion of 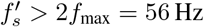.

Fig. 5 shows the power spectral density (PSD) estimate 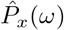 of a square wave test signal 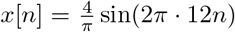, as well as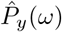, the PSD estimate of the filter output *y*[*n*]. The filter achieves negligible pass-band distortion with sharp transition-band roll-off and around 80 dB of attenuation in the stop-band.

### E. Decoding Algorithms

Consider an EEG signal tensor 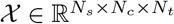 with *N*_*t*_ trials, each with *N*_*c*_ channels and *N*_*s*_ samples. In the context of CCA for SSVEP decoding with candidate frequencies *f*_*k*_ ∈ ℱ, observations from a given trial 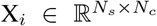 are compared with a sine-cosine reference Y_*k*_ generated as follows:

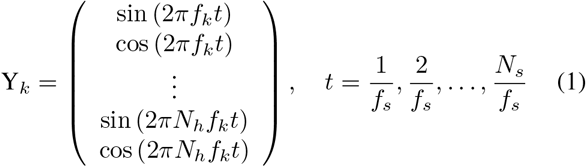

where *f*_*s*_ is the sampling frequency and *N*_*h*_ is the selected number of harmonics. Specifically, for each candidate frequency *f*_*k*_ ∈ℱ, CCA computes spatial filters (canonical weights) **w**_*X*_ and 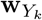 such that

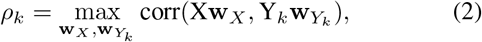

where *ρ*_*k*_ is the canonical correlation associated with candidate frequency *f*_*k*_ and corr(·) denotes Pearson correlation. The target frequency is selected as that which corresponds to the largest canonical correlation *ρ*_*k*_.

MsetCCA is an extension of standard CCA to multiple data sets or partitions introduced in [8]. Its objective is to maximise the correlation between canonical variables from many sets of observations for a given stimulus frequency *f*_*k*_. Although several objective functions for MsetCCA exist, the MAXVAR objective is explored in this work for its intuitive extension to ordinary CCA with multiple variable sets. Assuming all trial sets 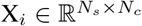 in are normalised to have zero mean and unit variance, the MAXVAR objective for candidate frequency *f*_*k*_ is defined as:

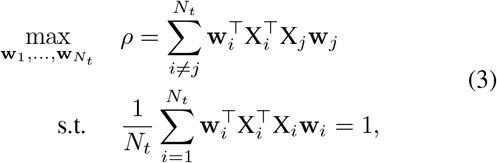

The expansion in [8] shows how (3) can be reformulated into a generalised eigenvalue problem. The optimal reference 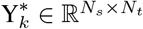 for *f*_*k*_ can be computed accordingly:

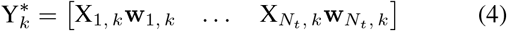

Following this calibration process, the standard CCA algorithm can be used for inference by comparing a test observation 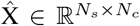 with the optimised reference set 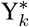 corresponding to each candidate frequency *f*_*k*_ instead of a data-independent harmonic reference signal. This important distinction is what allows MsetCCA to incorporate dynamics from historical data to improve SNR.

For GCCA, [9] proposed using historical observations (as in MsetCCA) in conjunction with a harmonic reference signal as in standard CCA to improve the algorithm’s robustness and reduce calibration requirements [9]. For brevity, the stimulus frequency index *k* is omitted in the following explanation since the procedure is identical and independent across stimulus frequencies *f*_*k*_ ∈ ℱ. Given *N*_*t*_ calibration trials, GCCA employs a template matrix 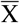 computed as the arithmetic mean across all trials: 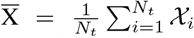 The concatenated signal matrix X^*c*^ is formed by unrolling *𝒳* along the trial axis: 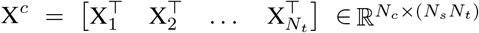. The template and sinusoidal reference matrices are concatenated similarly to match the dimensions of 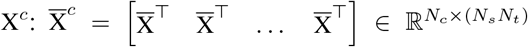 and 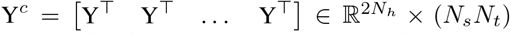 where Y follows the definition in (1). Consider the augmented spatial filter vector 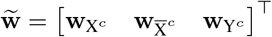 where, for example, component weight 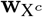 corresponds to concatenated signal matrix X^*c*^, and so on. Similarly, the augmented signal matrix is defined as 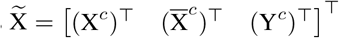. The objective of GCCA can then be expressed as

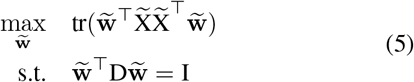

where D, the diagonal within-set block covariance matrix, is defined as

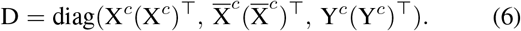

Using Lagrange duality as explored in [17], (5) can also be reformulated as a generalised eigenvalue problem of the form

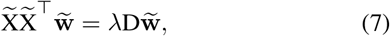

where the optimal combined weight 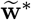 is the eigenvector corresponding to the largest eigenvalue *λ*^*^. Given a test observation 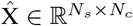, the three spatial filter components of 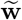 are used to compute two independent correlations: 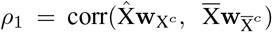 and 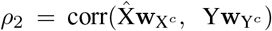. The combined output correlation is computed as 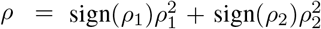. The target frequency is selected as 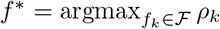.

### F. Firmware Implementation

All on-device firmware was implemented using MicroPython, a subset of the Python 3 standard library that is optimised for microcontrollers and other resource-constrained targets [18]. It is a full Python compiler implemented in C99 that runs natively on several architectures, e.g. x86, ARM and Xtensa, and is compact enough to run within 256 kB of ROM and 16 kB of RAM. Extensive use was made of open source MicroPython package ulab that provides a subset of the NumPy numerical computing library which offers high performance array computing and linear algebra functionality [19].

Each of the mentioned decoding algorithms can be formulated as a generalised eigenvalue problem in which the optimal weight/spatial filter can be found by solving for the eigenvector corresponding to the largest eigenvalue of the matrix of interest. A generalised eigenvalue solver was thus implemented in firmware for this purpose. Unfortunately, the eigenvalue solver offered by the ulab package can only be used on symmetric matrices. Eigenpairs for asymmetric matrices were estimated iteratively using a the QR iteration algorithm from [20].

An MQTT client was also implemented in firmware to allow the prototype to stream data over Wi-Fi to Amazon Web Services IoT Core for processing, storage and forwarding to a client-facing web interface. This enables multiple such BCI devices, each identified by a unique client ID, to stream data simultaneously.

### G. Data Acquisition

Due to COVID-19 limitations, all experimental data was recorded on the first author. The SSVEP stimulus squares interface shown in Fig. 4 were displayed on an 11” tablet with 120 Hz refresh rate. The screen was approximately 50 cm away from the subject’s face at 45 degrees below the horizontal plane through the subject’s eye line. Calibration of the BCI headset required the test subject to focus on a given stimulus square (as in Fig. 4) flashing at *f*_*k*_ Hz for *T* seconds. This process was repeated *p* times for each stimulus frequency *f*_*k*_ ∈ℱ, yielding a calibration signal tensor 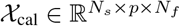 where *N*_*f*_ =| ℱ|= 3. The window length *T* corresponds to the number of samples *N*_*s*_ in each distinct trial. For example, a trial with *T* = 1 s corresponds to *N*_*s*_ = 256 samples at *f*_*s*_ = 256 Hz and 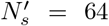 samples (as actually stored in memory) after downsampling to 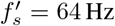. Experimental parameters *p* and *T* were varied to explore the impact of the number and length of calibration trials on out-of-sample decoding accuracy. Within each such experiment, *T* was kept constant across calibration and test trials.

**Fig. 4:**
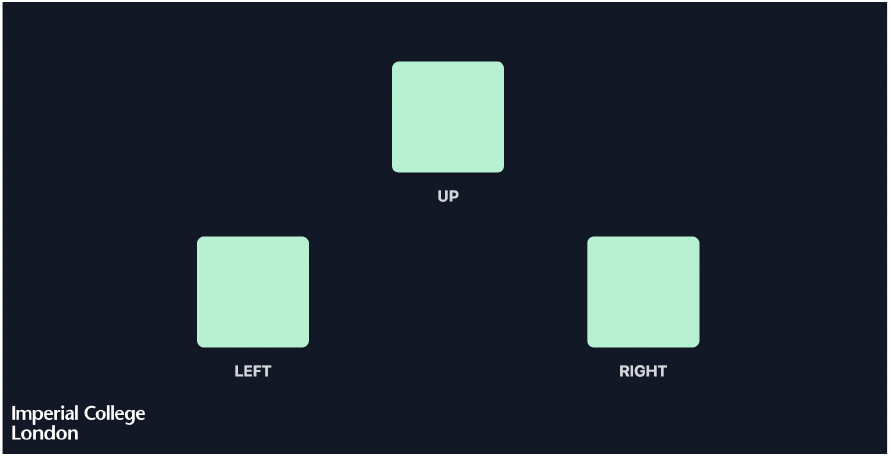
Web page for displaying SSVEP stimuli

**Fig. 5:**
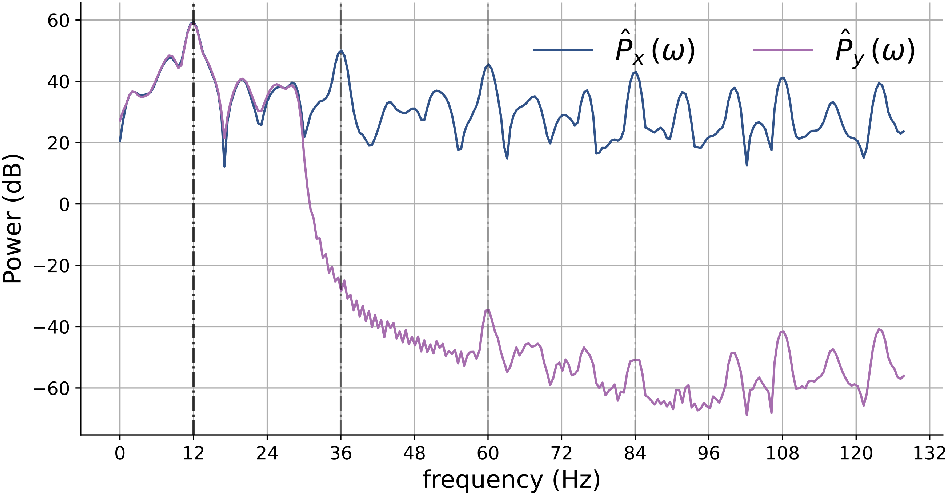
Power spectral density estimates 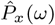 and 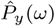 for a 12 Hz square wave input and its filtered output, respectively.

## III. Results

Leave-p-out cross-validation (LpO CV) was employed in evaluating test results to minimise the risk of selection bias and/or over-fitting. Typically, in this experiment, *n* = 7 trials were collected with the number of calibration trials varying in the range *p*∈ [1, 4], *p*∈ ℕ. For each performance metric, such as accuracy and ITR, reported results were obtained by computing the arithmetic mean of results from evaluating all combinations (CV folds) of calibrating on *p* trials and validating using the remaining *n* − *p* trials.

Fig. 6 shows the decoding accuracy for each stimulus frequency over a range of window lengths for the MsetCCA and GCCA algorithms. In each figure, two sets of results are displayed: one using *p* = 1 calibration trial for optimising spatial filters and another using *p* = 3 calibration trials prior to inference. All results shown represent cross-validated test accuracy evaluated on out-of-sample data.

**Fig. 6:**
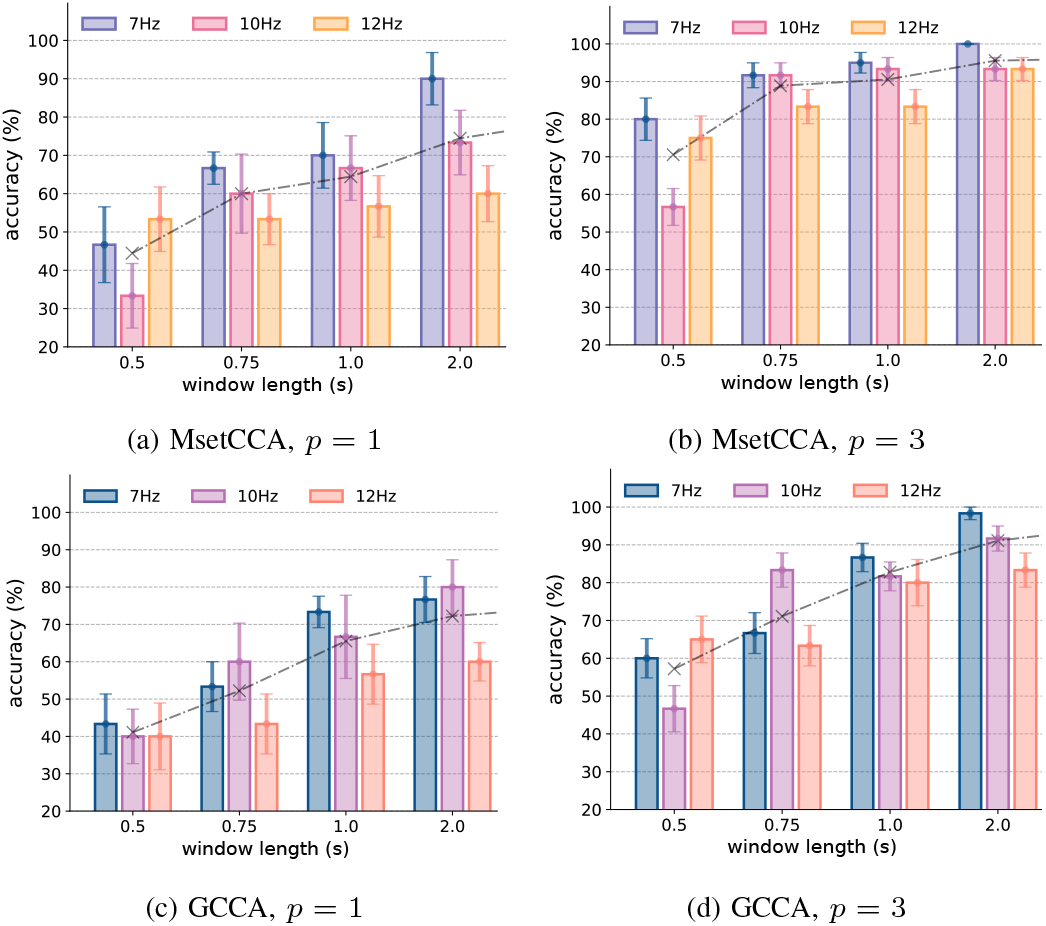
MsetCCA and GCCA decoding accuracies for varying trial length *T* and number of calibration trials *p*. Error bars denote standard error of the mean.

Table I gives a compilation of decoding performance metrics for the MsetCCA and GCCA algorithms. Results are reported over varying numbers of calibration trials *p* and recording window lengths *T*. Average accuracy 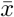 across frequencies and CV folds and corresponding standard error 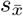, as well as ITR are given. ITR was calculated using the following expression:

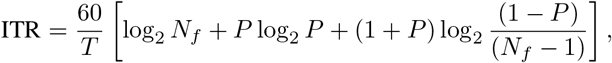

where *P* is decoding accuracy, *T* is the window length in seconds and *N*_*f*_ is the number of frequency targets. *P* = 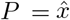 and *N*_*f*_ = 3 in this study. Notably, performing *p* = 4 calibration trials with *T* = 0.75 s yielded decoding accuracy of 95.6 *±* 3.74% with an ITR of 102.3 bits/min using the MsetCCA algorithm.

**TABLE I:**
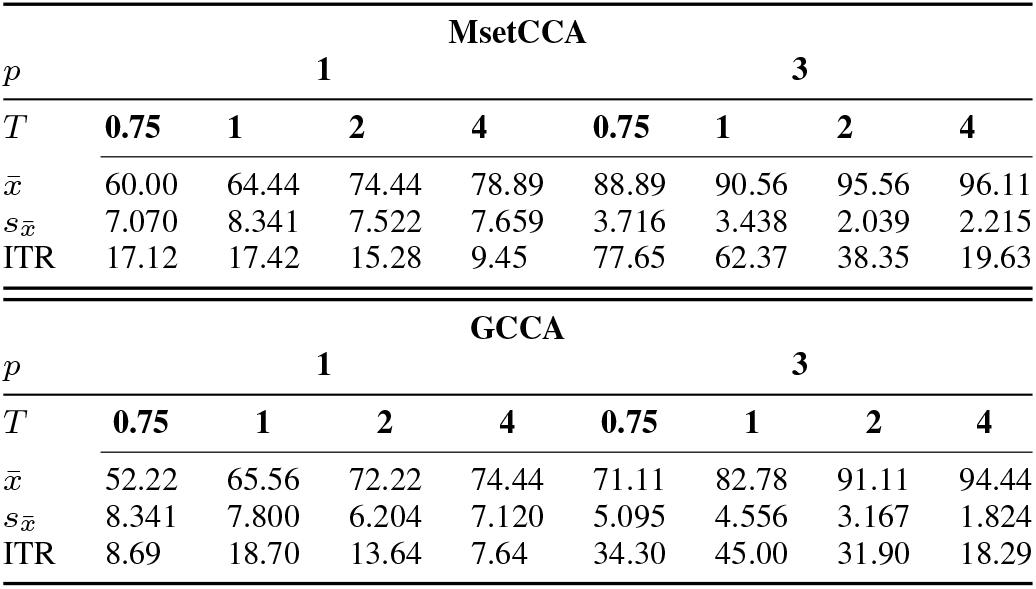
Summary of MsetCCA and GCCA decoding performance. 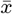 and 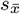 are given in %. ITR given in bits/min.

## IV. Discussion

The results obtained in this project suggest a very promising first prototype. It should be emphasised that these results were computed on data collected exclusively from the author. Several trials were conducted under various conditions and a stringent evaluation procedure was employed in an attempt to provide more robust results given this limitation.

For both algorithms, trends in average classification accuracy across frequencies show a strong positive correlation with *T* ; an expected result. However, increasing *T* leads to significantly reduced ITRs. The results suggest that this trade-off should be slightly more biased towards having more, shorter trials (larger *p*, decreased *T*). A possible explanation is that with larger *p*, calibration can be performed on more temporally-distinct trials which may capture a more robust representation of underlying EEG dynamics.

Decoding performance summaries in Table I clearly demonstrate that MsetCCA was superior to GCCA in this experiment. This may be as a result of being constrained to only use *N*_*h*_ = 1 harmonic in the harmonic reference signal compared to typical values of *N*_*h*_ = 3 or greater in similar studies without the same memory and computational constraints.

Of the devices in the EEG-SSVEP decoding literature, the easiest comparison of this work is to Acampora et al. in [14]. Whereas [14] decoded four SSVEP frequencies with a single channel EEG device, this work decoded three SSVEP frequencies with the same. Similar decoding accuracies were obtained for similar time windows, although the system in [14] was trained on 563 s of data and tested on 141 s. As such, while the system in this work is somewhat less capable due to it decoding fewer frequencies, it is significantly cheaper.

It was found that when performing calibration and subsequent inference on data collected during different sessions, performance degraded significantly. Here, distinct sessions are distinguished by the user removing the BCI headset and reinstalling. This degradation in performance was likely caused by lack of standardised positioning of electrodes; both in terms of spatial location and contact quality. This challenge is likely exacerbated by only having a single active EEG channel. As such, the device should be re-calibrated if its position is adjusted on the subject.

Future work should explore a more robust mechanism for achieving consistent flicker frequency of SSVEP stimuli. While basic animation using vanilla Javascript proved sufficient when using a tablet with a display refresh rate of up to 120Hz, this approach may yield inconsistent performance on mobile or edge display devices. WebGL could offer a suitable alternative.

## V. Conclusion

The findings suggest that a fully functional, if basic, BCI device can be deployed in a small, self-contained package that is both completely mobile and extremely cost-effective. While likely slightly less capable than existing commercial counterparts, the device investigated in this project poses an interesting low-cost option for large scale EEG data acquisition, testing and experimentation. It could also become a valuable and accessible tool for engaging and educating young audiences about neurophysiology and neurotechnology.

## Notes

### Competing Interest Statement

The authors have declared no competing interest.

